# Single-Objective Evanescent Scattering Microscopy for Imaging Single Proteins and Binding Kinetics

**DOI:** 10.1101/2022.02.04.479201

**Authors:** Pengfei Zhang, Rui Wang, Zijian Wan, Xinyu Zhou, Guangzhong Ma, Jayeeta Kolay, Jiapei Jiang, Shaopeng Wang

## Abstract

Plasmonic scattering microscopy has advanced the evanescent detection approaches by offering wide-field single-molecule imaging capability. However, two limitations prevent the broader application of plasmonic single-molecule imaging. One is the heating effect accompanying the plasmonic enhancement, and the other is the complicated system structure resulting from the two-objective optical arrangement. Here, we report single-objective evanescent scattering microscopy. The evanescent field is created by total internal reflection instead of the surface plasmon resonance on the gold film. As a result, the sensing substrate without gold film produces little heat, and allows excitation and observation using one objective. In addition, this system enables quantification of protein binding kinetics by simultaneously counting the binding of individual molecules and recording their binding sites with nanometer precision, providing a digital method to measure binding kinetics with high spatiotemporal resolution. This work may pave a road for label-free single protein analysis in conventional microscopy.

**Teaser:** Label-free single-molecule imaging on a total internal reflection fluorescence objective.

## Introduction

Determining molecular interactions is vital for drug screening, molecular diagnosis, biochemical analysis, and understanding the biological processes at the molecular level(*1, 2*). To meet this need, evanescent detection approaches have been developed as a powerful tool for detecting molecules and quantifying the molecular interaction dynamics. These approaches are accomplished with different processes, such as total internal reflection (TIR)(*3*), surface plasmon resonance (SPR)(*4*), optical waveguide(*5, 6*), manipulating Bloch surface waves(*7*), optical microcavities(*8*), and plasmonic resonators(*9*). The shared feature of these processes is creating an evanescent field localized within several hundred nanometers from the sensor surface for measurement. The evanescent field has exponentially decaying intensity in the axial direction, thus allowing tracking analyte motion with resolution down to sub-nanometer level by monitoring signal intensity variations in real-time(*10-14*). More importantly, the evanescent field can significantly reduce the illumination volume for enhancing light-analyte interaction and reducing the environmental noise, which is responsible for the high sensitivity of evanescent detection(*15*). As a result, the combination of evanescent detection and fluorescence approaches, namely the total internal reflection fluorescence, has become one of the most popular single-molecule detection schemes in the last decades(*16*). In recent years, the label-free evanescent single-molecule sensing approaches were intensely studied to analyze intrinsic molecular properties such as mass and quantify molecular interaction kinetics without labels(*9, 10, 17-19*). Recently developed plasmonic scattering microscopy (PSM) has further advanced this field by providing single-molecule imaging capability for parallelly monitoring the molecular interaction process in different locations(*20-22*). However, two factors prevent the broader applications of PSM. One is the heating effect accompanying the plasmonic enhancement(*23*), limiting its applications for detecting fragile biological molecules under normal conditions(*21*); the other is the complicated optical system structure as well as the strictly sealed flow cell for clear observations with the second objective. Specifically, the flow channel needs to be constructed with a thin cover glass as top window and a channel height less than 100 micrometers for correctable image distortion(*24*), which makes it challenging to combine with other technologies, such as electrochemical microscopy(*25-28*), to analyze additional molecular properties such as charge and charge-to-mass ratio.

Here, we show that label-free evanescent single-molecule imaging can be achieved on a single total internal reflection fluorescence objective. TIR replaces SPR to provide the evanescent illumination for single-molecule imaging. As a result, no gold film is coated on the sensing substrate, thus allowing excitation and observation using one objective without heating effect. We also show that single protein molecules can be detected and identified based on their mass and specific binding to the corresponding antibodies. Additionally, we demonstrate measurement of protein binding kinetics, which is one mainstream application of evanescent detection approaches, by digitally counting the binding and unbinding of individual molecules. Furthermore, we also show that this system can record the protein binding sites with nanometer precision, providing a solution to analyze molecular interaction with high spatiotemporal resolution.

## Results

### Imaging principles

We excite evanescent waves using TIR configuration by directing light beyond critical angle via an oil-immersion objective onto an Indium-Tin-Oxide (ITO) coated coverslip placed on the objective (Fig. 1, and Fig. S1 for details). The ITO film was employed to provide nanometer-scale surface roughness profiles instead of the gold film used in the PSM (Fig. S2). As both the ITO film and coverslip have good optical transparency, we can collect the evanescent waves scattered by the ITO surface and proteins with the same objective. To reduce the optical energy loss in the path, the polarization beam splitting strategy is used here(*29, 30*). As shown in Fig. 1A, the *s*-polarized incident beam becomes circularly polarized after passing the quarter-wave plate. Then, the circularly polarized reflection beam and evanescent waves scattered by the ITO surface and proteins become *p*-polarized after passing the quarter-wave plate and then are reflected to the detection path by a polarization beam splitter. Finally, one blocker with a diameter of 4 mm was applied to block the intense reflection beam. Thus, we can clearly see the background (*E*_*b*_) created by evanescent waves scattered by the surface roughness of ITO film (Fig. 1B). The *E*_*b*_ can interfere with the evanescent waves scattered by the proteins (*E*_*s*_). Consequently, the image is given by

**Fig. 1.**
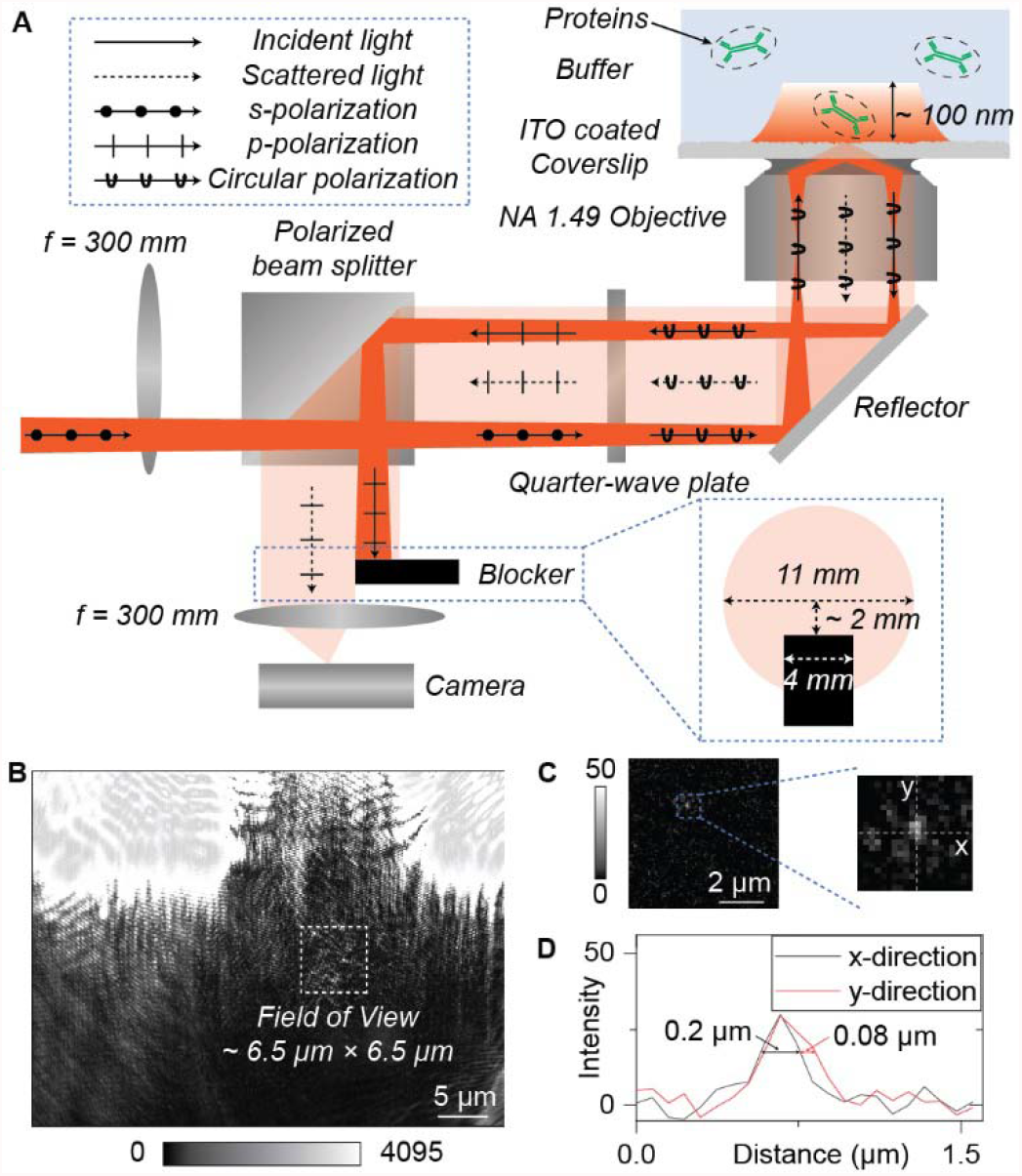
Setup and principle of single-objective ESM. **A;** Simplified sketch of the optical setup, where evanescent waves are excited on the surface of ITO coated coverslip using total internal reflection configuration. Scattering of the evanescent waves by a protein (*E*_*s*_) and by the ITO surface (*E*_*b*_) is collected from the bottom. One blocker with a diameter of 4 mm is employed to stop the reflection beam to avoid overexposure. A detailed description of the setup can be found in Fig S1. **B**; Raw image captured by the camera. The system field of view was marked with a dashed square. **C;** ESM image of one IgG protein molecule after removing *E*_*b*_. **D**; Profiles of marked lines in **C**. Incident light intensity, 150□kW□cm^-2^; camera exposure time, 0.5□ms.

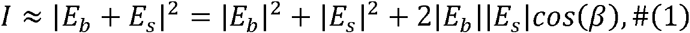

where *β* is the phase difference between light scattered by the ITO surface and proteins. The *E*_*b*_ created by the high-density ITO islands on the surface is much larger than *E*_*s*_ created by proteins, and thus the interference term, 2|*E*_*b*_| |*E*_*s*_| cos(*β*), dominates the sensor output and produces the image contrast that scales with protein volume, or the molecular weight. To differentiate this plasmonic imaging method from PSM, we refer it to as evanescent scattering microscopy (ESM). The ESM does not rely on the gold film to produce an evanescent wave and therefore permits the incident intensity of 150 kW/cm^2^ used here, which is 50 times higher than the PSM(*20*), making it possible to image smaller proteins (Fig. S3).

To achieve the high contrast ESM image of single proteins, it is necessary to remove |*E*_*b*_|^2^ in equation (1), which is achieved with the following imaging processing algorithm. Starting from the images captured with a high frame rate, usually 2000 frames per second, we first average the image frames within usually 20 ms to suppress random noise in the images (Fig. S4). We then obtain differential images by subtracting a previous frame from each frame. The subtraction removes background features and captures the binding of a protein to the surface. The codes for data processing have been published previously(*14, 20, 21*). Fig. 1C shows an ESM image of one immunoglobulin G (IgG) molecule, where the point spread function is wider in vertical than horizontal direction (Fig. 1D). This should be because the blocker prevents high spatial frequency components from passing, leading to smaller effective numerical aperture and thus worse spatial resolution in the vertical direction. As a result, we determined the image intensity by integrating the intensities of all pixels within the disk centered around the protein’s binding position with the diameter selected from the length of the axis of the point spread function in the vertical direction.

### Detection of single proteins

We validated single-objective ESM by imaging the proteins with different molecular weights, including bovine serum albumin (BSA, 66 kDa), goat IgG (150 kDa), human immunoglobulin A (IgA, 385 kDa), and human immunoglobulin M (IgM, 950 kDa). For each molecular weight, the proteins dispersed in phosphate-buffered saline (PBS) buffer were introduced into a solution well mounted on the ITO surface, and the binding of the proteins to the surface was recorded over time. The ITO surface was modified with N-hydroxysuccinimide (NHS) groups to increase the binding rate (Methods). Fig. 2A shows the ESM images of BSA, IgG, IgA, and IgM proteins binding on the surface, and the ESM image achieved by flowing PBS buffer was also shown as control. The ESM images reveal the individual proteins as bright spots due to the constructive interference between the evanescent waves scattered by the ITO surface and proteins.(*31-33*), and the number in the intensity bar indicates the grayscale intensity range. We tracked and counted the individual protein binding events over 10□min and constructed an image intensity histogram from the multiple protein molecules for each molecular weight. The image intensity histogram of the individual proteins follows a Gaussian distribution (Fig. 2B), and ESM image contrast scales with the molecular weight of proteins, which is clearly shown by the intensity histograms (Fig. 2C). By fitting the histograms with a Gaussian distribution, the mean intensity of each protein was extracted and presented in Fig. 2B. Plotting the mean image intensity versus molecular weight reveals a linear relationship (Fig. 2D), confirming that the ESM image intensity is a measure of protein volume and therefore the molecular mass. The signal-to-noise ratio (SNR) for ESM imaging of BSA proteins is estimated to be ∼10 (Section S1), while the PSM usually can only achieve similar SNR for imaging IgA(*20*), which is ∼6 times heavier than BSA, demonstrating the advantages of single-objective ESM for single protein analysis.

**Fig. 2.**
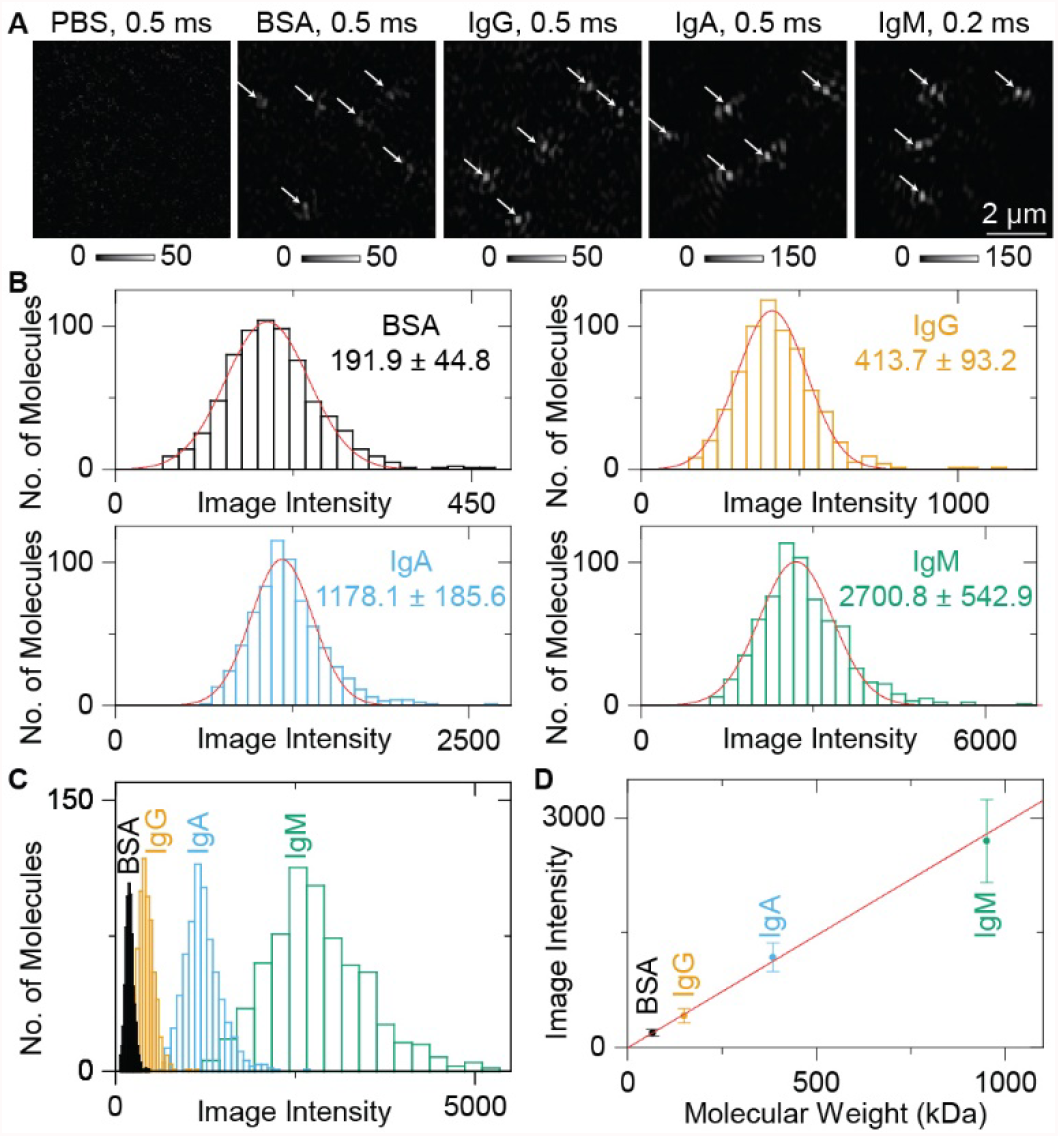
Imaging single proteins with single-objective ESM. **A;** ESM imaging of BSA, IgG, IgA, and IgM protein molecules binding on the surface. The white arrows indicate the binding sites. The ESM image achieved by flowing PBS buffer was presented as control. **B**; Image intensity histograms, where the solid lines are Gaussian fittings, of BSA, IgG, IgA, and IgM proteins by counting the individual protein binding events. **C**; Image intensity histograms of BSA, IgG, IgA, and IgM proteins in the same scale. **D**; ESM image intensity versus molecular weight. The image intensity at the center of error bar for each molecular weight is the mean value of the corresponding histograms in **B**. The error bars indicate the full width at half maximum of the Gaussian fitting of corresponding histograms. Incident light intensity, 150□kW□cm−^2^. Camera exposure time is 0.5□ms for BSA, IgG, and IgA, and 0.2ms for IgM.

### Identification of single proteins

To identify single proteins with single-objective ESM, we coated the ITO surface with anti-BSA and studied the specific binding of BSA to anti-BSA (Fig. 3A). On exposure to BSA, the binding of single BSA molecules to anti-BSA took place immediately, which was observed as the bright spots appearing one at a time on the surface. Movie S1 reveals the dynamic binding process, and Fig. 3A shows a few snapshots. To view all the BSA binding events on the surface on the Nth frame, we integrate the differential images from 1 to N as described before(*20*). We counted the individual BSA binding events over 10□min and constructed a molecular weight histogram using the calibration curve shown in Fig. 2D, showing a major peak due to single BSA molecules (Fig. 3B). As a negative control experiment, we flowed IgA over the anti-BSA coated ITO surface. Unlike the case of BSA, where a bright spot appears and stays on the surface, bright spots (IgA proteins) show up on the surface only transiently (Movie S1 and Fig. 3C), which is expected because IgA does not bind specifically to anti-BSA. For the negative control experiment, the average period is set to be 1.5 ms for improving temporal resolution to avoid motion blur. The identification experiment confirms that the single-objective ESM can image specific single proteins based on affinity recognition.

**Fig. 3.**
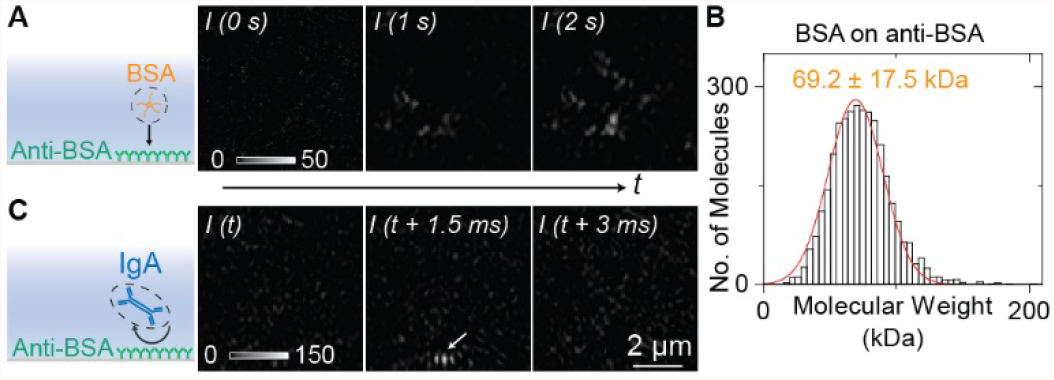
Identifying single proteins with single-objective ESM. **A;** Integrated ESM snapshots showing binding of BSA to anti-BSA immobilized on the surface. **B**; Molecular weight histogram of BSA molecules, where the solid lines are Gaussian fitting. **C**; Negative control experiment, exposing of IgA to anti-BSA surface. The white arrow indicates the IgA hitting event on the surface. Incident light intensity, 150□kW□cm^-2^; camera exposure time, 0.5□ms.

### Quantification of protein binding kinetics

The most powerful application of PSM is measuring the molecular binding directly by digital counting for binding kinetics analysis, rather than resonant angle shift on the ensemble SPR. The latter requires further correction on the refractive index of the solution for accurate binding kinetics measurement(*20, 21*). We show here that the single-objective ESM can also measure the protein binding kinetics at the single-molecule level by counting the binding and unbinding of single molecules. As a proof-of-concept, we studied IgA binding to anti-IgA (Fig. 4A). We first flowed IgA over an anti-IgA coated ITO surface to monitor the association process, then flowed PBS buffer over the sensor surface to dissociate IgA from anti-IgA. We tracked the association and dissociation processes by counting the individual IgA molecules in real-time. Plotting the number of bound IgA versus time produces binding kinetics curves. Fitting of the curves with the first-order binding kinetics model determines the association (*k*_*on*_) and dissociation (*k*_*off*_) rate constants, which are 2.3□×□10^5^□M^−1^□s^−1^ and 2.0□×□10^−4^□s^−1^, respectively. From *k*_*on*_ and *k*_*off*_, the equilibrium dissociation constant (*K*_*D*_□=□*k*_*off*_/*k*_*on*_) is determined to be 870□pM. These values agree with the published results(*20, 21*). In addition, the mean intensity changes associated with the binding and unbinding of events are consistent with the molecular weight of an IgA molecule, confirming the detection of single IgA proteins (Fig. 4B).

**Fig. 4.**
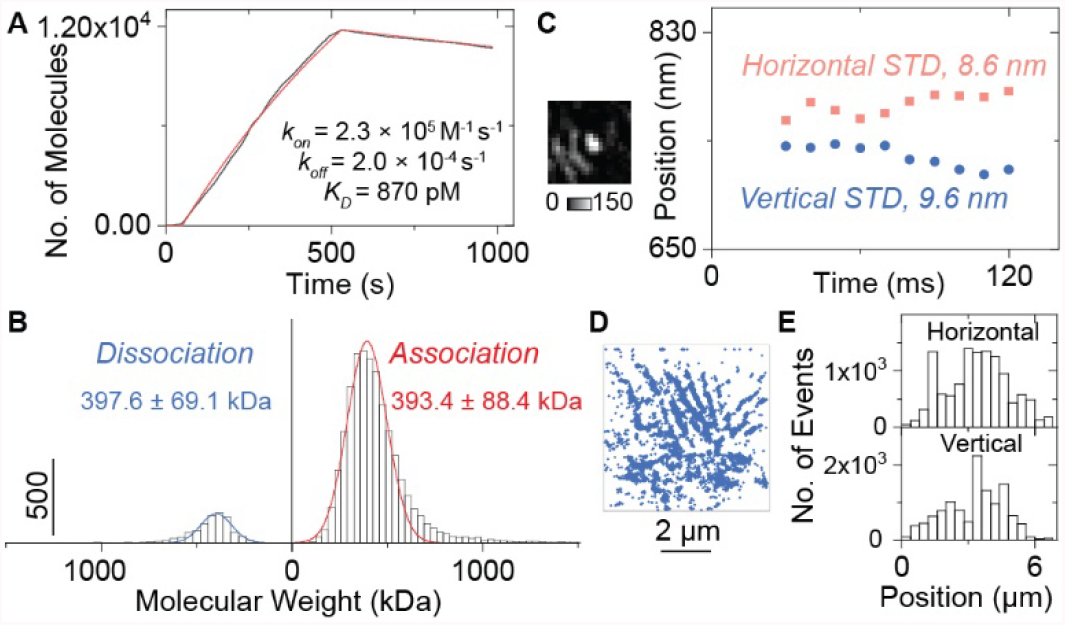
Single-molecule measurement of binding kinetics with single-objective ESM. **A;** Kinetics of 5 nM IgA binding to anti-IgA determined by digitally counting the binding/unbinding of single molecules. **B;** Molecular weight histograms of binding and unbinding of individual IgA molecules, where the solid lines are Gaussian fittings. The mean values of the histograms are presented in the figure. **C;** One snapshot of IgA binding on the surface, and super-resolution localization determined by two-dimensional fitting in the horizontal and vertical directions. The localization precision is estimated with the standard deviation (STD) of the IgA binding positions determined at different times. **D**; Super-resolution image, showing the localized positions of the individual IgA binding to anti-IgA measured in **A. E**; Statistics of localized positions of the individual IgA binding to anti-IgA in horizontal and vertical directions shown in **D**. Incident light intensity, 150□kW□cm^-2^; camera exposure time, 0.5□ms.

Besides the high temporal resolution for binding kinetics analysis at the single-molecule level, the single-objective ESM also permits the super localization of protein binding sites with high precision owing to that it employs an oil immersion objective with a numerical aperture of 1.49, which is ∼3.5 times higher than the observation objective used in PSM(*20*). Two-dimensional Gaussian fitting was employed to localize the protein binding sites(*34*). We track one IgA protein binding on the ITO surface for 100 ms with a temporal resolution of 10 ms to estimate the localization error (Movie S2 and Fig. 4C). The localization precision is estimated to be 8.6 nm, and 9.6 nm at horizontal, and vertical directions, respectively. The lower localization precision in the vertical direction should be because of the smaller effective numerical aperture resulting from the reflection beam blocker. The localization precision in both directions is lower than theoretical predictions (Section S2), which should be because the tracking period of 100 ms is longer than the shot noise limited duration of ∼20 ms (Fig. S4). Then, the sites of IgA binding to anti-IgA measured in Fig. 4A were localized as shown in Fig. 4D. The binding probability is not uniform over the surface, which can be more clearly revealed by the statistical analysis in horizontal and vertical directions (Fig. 4E). The IgA prefer binding to a few areas over others on the ITO surface, which should be because the thin APTES film used for modifying ITO surface anti-IgA is usually not uniform due to the multilayer-island growth kinetics, resulting in nonuniform antibody coverage(*35*).

## Discussion

The image contrast of single-objective ESM arises from the interference of evanescent waves scattered by sensor surface and proteins, which is analogous to PSM. Single-objective ESM shares the advantage of PSM, namely that the evanescent enhancement effect enables comparable signal-to-noise ratio with either lower incident light power or wider field of view than conventional nonevanescent approaches. Compared to PSM, single-objective ESM shows two advantages. First, the ITO surface produces little heat, thus allowing the incident light intensity of up to 150 kW cm^-2^, which is ∼50 times higher than the PSM. This makes it possible to use single-objective ESM for reliably imaging medium-sized single proteins as small as the BSA (66 kDa), which is usually considered as the practical measurement limit for label-free single-molecule detection technologies(*10*). Second, the ITO coated coverslips have good transparency, thus allowing the excitation and detection with a single total internal reflection fluorescence objective, which has a much higher numerical aperture than the dry objective used for PSM(*20*). Owing to this, the single-objective ESM allows both higher collection efficiency and super-resolution localization precision. Furthermore, the single objective optical arrangement simplifies the system structure by removing the second objective on the top of the sensor surface, thus eliminating the image distortion resulting from the refraction of the water layer and allowing the applications of conventional flow cells, rather than the complicated flow cell with a thin cover glass as a top window for PSM. The open top optical arrangement also makes it easy to combine with other technologies, such as electrochemical microscopy, to analyze additional molecular properties such as charge.

We have demonstrated the evanescent single-molecule imaging with TIR configuration on a single-objective ESM system. The light scattered by the ITO surface is used as a reference for interfering with light scattered by proteins, providing the quantitative mass imaging of single proteins. We also show that this approach can identify single proteins and quantify the protein binding kinetics by digitally counting the binding and unbinding of single proteins. In addition, the single-objective ESM can also provide super-resolution localizations of protein binding sites, thus providing one solution for label-free mapping of target proteins on the bio membrane by dynamically monitoring the antibody binding. The single-objective ESM also presents a solution to achieve label-free single protein imaging with open top inverted microscopy configuration, which can be easily combined with other technologies. Thus, we anticipate that the single-objective ESM can become an essential tool for analyzing single molecule behaviors, especially when combined with electrochemical microscopic imaging for a systematic understanding of protein distributions and activities on the biological surfaces.

## Materials and Methods

### Materials

The No.1 ITO Coated Cover Slips (22×22 mm, Catalog No. 06470-AB) were purchased from SPI Supplies (West Chester, PA, US). Isopropyl alcohol (IPA, Catalog No. BDH2032-1GLP) was purchased from VWR (Radnor, PA, US). Hydrogen peroxide (H_2_O_2_, 30%, Catalog No. H1009), (3-Aminopropyl)triethoxysilane (APTES, Catalog No. 440140), succinic anhydride (Catalog No. 239690), and bovine serum albumin (BSA, Catalog No. A7638), sodium hydroxide (NaOH, Catalog No. S5881) were purchased from Sigma-Aldrich (St. Louis, MO, US). Ammonium hydroxide (NH_3_H_2_O, 28.0 to 30.0%, Catalog No. C5103500-2.5D) was purchased from Mallinckrodt Reagents (Belmont, NC, US). 1-ethyl-3-(3-dimethylaminopropyl)carbodiimide hydrochloride (EDC, Catalog No. 22980) and Sulfo-NHS (N-hydroxysulfosuccinimide, Catalog No. 24510) were purchased from Thermo Scientific (Waltham, MA, US). Phosphate-Buffered Saline (PBS, Catalog No. 21-040-CV) was purchased from Corning (Corning, NY, US) and filtered with 0.22-µm filters (Millex-GS, Catalog No. SLGSM33SS) from Sigma-Aldrich (St. Louis, MO, US). Goat anti-human IgA (IgG, Catalog No. STAR141) was purchased from BIO-RAD (Hercules, CA, US). Human colostrum IgA (Catalog No. 16-13-090701) and Human IgM (Catalog No. 16-16-090713) were purchased from Athens Research and Technology (Athens, GA, US). Deionized water with resistivity of 18.2□MΩ□cm^−1^ was filtrated with a 0.22-µm filter and used in all experiments.

### Experimental setup

A 120-mW laser diode (L660P120, Thorlabs, Newton, NJ, US) is used as the light source to provide the incident wavelength of 660 nm. The laser diode is attached to a temperature-controlled mount (LDM56, Thorlabs), which is driven by a benchtop diode current controller (LDC205C, Thorlabs) and a temperature controller (TED200C, Thorlabs). Light from the laser diode is conditioned by an achromatic doublet lens group, and then focused to the back focal plane of a ×60 objective (Olympus APO N 60x Oil TIRF, NA 1.49) by a tube lens with focal length of 300□mm. The incident angle was adjusted by a manual translation stage to reach total internal reflection condition at 65° (XR25P-K2, Thorlabs). A combination of polarization beam splitter with quarter-wave plate is employed to separate the signal light from incident light. Reflection beams is blocked by a M4 screw with a diameter of 4 mm. The evanescent waves scattered by the ITO surface and proteins are collected by a camera (MQ003MG-CM, XIMEA). A detailed schematic representation of the optics can be found in Fig. S1.

### Surface functionalization

For measuring nonspecific binding of single proteins, the ITO coated coverslips were modified with active carboxyl groups using following steps. 1) The ITO coated coverslips were cleaned in the boiling solution mixing the NH_3_H_2_O, H_2_O_2_, and water with volume ratio of 1:1:5 for 1 hour to obtain clean hydroxylated ITO surfaces, where dropping water became a layer. 2) The ITO coated coverslips and container were washed twice with water, and then ultrasonically cleaned 2 times with water, and blew dry with nitrogen. 3) The hydroxylated ITO coated coverslips were incubated in boiling 1% APTES diluted with IPA for 3 hours to functionalize the surface with primary amine group. After processing, both solution and ITO coated coverslips should be clear if drying in second step is performed properly. 4) ITO coated coverslips and container were washed twice with IPA, ultrasonically cleaned twice with IPA, and blew dry with nitrogen. 5) Incubate the amino group modified ITO coated coverslips in 10 g L^-1^ succinic anhydride in water for 1 hour to obtain carboxylic group functionalized ITO coated coverslips. The pH of succinic anhydride solution is adjusted to 7.5∼8 with 1 M NaOH solution. The ITO coated coverslips and container were washed twice with water, ultrasonically cleaned twice with water, and then stored in the water prior to use. In the experiment, the surface was incubated in 40g L^-1^ EDC mixed with 11g L^-1^ Sulfo-NHS for 15□min to activate the carboxyl groups for connecting proteins. The EDC/NHS solution was filtered by a 0.22-µm filter before use. For specific binding kinetic analysis, the activated carboxylic group modified ITO coated coverslips were rinsed with PBS buffer, and then 20□nM anti-IgA in PBS buffer was applied to the surface and incubated for 1 hour to allow immobilization. Next, the surface was incubated in 1□mg□ml^−1^ BSA for 10□min to block any remaining activated carboxylic group to minimize nonspecific binding. Finally, the protein solution was flowed onto the surface for specific binding measurement.

### Data processing

The raw image sequence captured at high frame rate (2000□fps) was converted to an averaged-image sequence, by usually averaging images over every 20□ms using previously reported MATLAB program or the real-time averaging function of the camera recording software (XIMEA CamTool), to suppress shot noise. To remove the background, a differential image sequence was obtained by subtracting the previous frame using the Image Calculator plugin in ImageJ. The TrackMate plugin in ImageJ was employed to find and count particles or molecules. The ESM intensity of a particle or molecule was determined by integrating the intensities of all pixels within the disk centered around the protein’s binding position with the diameter selected from the length of the axis of the point spread function in the vertical direction. Origin 2019 was used to create data plots and histograms. Scrubber v.2.0a was used to determine the association and dissociation rate constants by fitting the curves in Fig. 4A with the first-order binding kinetics model.

## Author contributions

P. Z. and S. W. designed the research; P. Z. built the setup and performed the measurement and data analysis; R. W., X.Z., J. K., and J. J. contributed to the surface modification; Z. W. contributed to the AFM imaging; G. M. contributed to the data analysis; S. W. supervised the experiments; P. Z., R. W., and S. W. wrote the paper.

## Data and materials availability

The data for the reported analyses are available upon request from the corresponding author.

## Supplementary Materials

The PDF file includes:

Fig. S1. Optical setup for single-objective evanescent scattering microscopy.

Fig. S2. Surface roughness profiles of ITO surface.

Fig. S3. Evaluation of heating effect. Fig. S4. Estimation of shot noise limit.

Section S1. Theoretical prediction of signal-to-noise ratio.

Section S2. Estimation of super resolution localization.

Other Supplementary Materials for this manuscript include the following

Movie S1.

Dynamic binding of BSA to anti-BSA and negative control experiment of flowing IgA onto anti-BSA.

Movie S2.

Tracking of one IgA protein binding on the surface.

## References

1. M. Polanski, N. L. Anderson, A List of Candidate Cancer Biomarkers for Targeted Proteomics. Biomarker Insights 1, 1–48 (2006).

2. R. Santos et al., A comprehensive map of molecular drug targets. Nature Reviews Drug Discovery 16, 19–34 (2017).

3. R. M. Dickson, D. J. Norris, Y.-L. Tzeng, W. E. Moerner, Three-Dimensional Imaging of Single Molecules Solvated in Pores of Poly(acrylamide) Gels. Science 274, 966–968 (1996).

4. J. Homola, Surface Plasmon Resonance Sensors for Detection of Chemical and Biological Species. Chemical Reviews 108, 462–493 (2008).

5. M. Sjöberg et al., Time-Resolved and Label-Free Evanescent Light-Scattering Microscopy for Mass Quantification of Protein Binding to Single Lipid Vesicles. Nano Letters 21, 4622–4628 (2021).

6. T.-H. Lee, D. J. Hirst, K. Kulkarni, M. P. Del Borgo, M.-I. Aguilar, Exploring Molecular-Biomembrane Interactions with Surface Plasmon Resonance and Dual Polarization Interferometry Technology: Expanding the Spotlight onto Biomembrane Structure. Chemical Reviews 118, 5392–5487 (2018).

7. Y. Kuai et al., Label-free surface-sensitive photonic microscopy with high spatial resolution using azimuthal rotation illumination. Science Advances 5, eaav5335 (2019).

8. M. D. Baaske, M. R. Foreman, F. Vollmer, Single-molecule nucleic acid interactions monitored on a label-free microcavity biosensor platform. Nature Nanotechnology 9, 933–939 (2014).

9. M. D. Baaske, N. Asgari, D. Punj, M. Orrit, Nanosecond time scale transient optoplasmonic detection of single proteins. Science Advances 8, eabl5576 (2022).

10. N. P. Mauranyapin, L. S. Madsen, M. A. Taylor, M. Waleed, W. P. Bowen, Evanescent single-molecule biosensing with quantum-limited precision. Nature Photonics 11, 477–481 (2017).

11. Z. Chen et al., Light-Driven Nano-oscillators for Label-Free Single-Molecule Monitoring of MicroRNA. Nano Letters 18, 3759–3765 (2018).

12. H. Wang, Z. Tang, Y. Wang, G. Ma, N. Tao, Probing Single Molecule Binding and Free Energy Profile with Plasmonic Imaging of Nanoparticles. Journal of the American Chemical Society 141, 16071–16078 (2019).

13. G. Ma, Z. Wan, Y. Yang, W. Jing, S. Wang, Three-Dimensional Tracking of Tethered Particles for Probing Nanometer-Scale Single-Molecule Dynamics Using a Plasmonic Microscope. ACS Sensors 6, 4234–4243 (2021).

14. P. Zhang et al., Label-Free Imaging of Nanoscale Displacements and Free-Energy Profiles of Focal Adhesions with Plasmonic Scattering Microscopy. ACS Sensors 6, 4244–4254 (2021).

15. J. O. Arroyo, P. Kukura, Non-fluorescent schemes for single-molecule detection, imaging and spectroscopy. Nature Photonics 10, 11–17 (2016).

16. A. Daniel, Selective imaging of surface fluorescence with very high aperture microscope objectives. Journal of Biomedical Optics 6, 6–13 (2001).

17. P. Zijlstra, P. M. R. Paulo, M. Orrit, Optical detection of single non-absorbing molecules using the surface plasmon resonance of a gold nanorod. Nature Nanotechnology 7, 379–382 (2012).

18. Y. Pang, R. Gordon, Optical Trapping of a Single Protein. Nano Letters 12, 402–406 (2012).

19. E. Kim, M. D. Baaske, I. Schuldes, P. S. Wilsch, F. Vollmer, Label-free optical detection of single enzyme-reactant reactions and associated conformational changes. Science Advances 3, e1603044 (2017).

20. P. Zhang et al., Plasmonic scattering imaging of single proteins and binding kinetics. Nature Methods 17, 1010–1017 (2020).

21. P. Zhang, G. Ma, Z. Wan, S. Wang, Quantification of Single-Molecule Protein Binding Kinetics in Complex Media with Prism-Coupled Plasmonic Scattering Imaging. ACS Sensors 6, 1357–1366 (2021).

22. P. Zhang, S. Wang, Real-Time analysis of exosome secretion of single cells with single molecule imaging. Biocell 45, 1449–1451 (2021).

23. M. L. Brongersma, N. J. Halas, P. Nordlander, Plasmon-induced hot carrier science and technology. Nature Nanotechnology 10, 25–34 (2015).

24. M. L. Martin-Fernandez, C. J. Tynan, S. E. D. Webb, A ‘pocket guide’ to total internal reflection fluorescence. Journal of Microscopy 252, 16–22 (2013).

25. G. Ma et al., Optical imaging of single-protein size, charge, mobility, and binding. Nature Communications 11, 4768 (2020).

26. X. Shan, U. Patel, S. Wang, R. Iglesias, N. Tao, Imaging Local Electrochemical Current via Surface Plasmon Resonance. Science 327, 1363–1366 (2010).

27. X. Shan et al., Imaging the electrocatalytic activity of single nanoparticles. Nature Nanotechnology 7, 668–672 (2012).

28. W. Wang et al., Single cells and intracellular processes studied by a plasmonic-based electrochemical impedance microscopy. Nature Chemistry 3, 249–255 (2011).

29. D. Cole, G. Young, A. Weigel, A. Sebesta, P. Kukura, Label-Free Single-Molecule Imaging with Numerical-Aperture-Shaped Interferometric Scattering Microscopy. ACS Photonics 4, 211–216 (2017).

30. G. Young et al., Quantitative mass imaging of single biological macromolecules. Science 360, 423–427 (2018).

31. J. Berk, M. R. Foreman, Role of multiple scattering in single particle perturbations in absorbing random media. Physical Review Research 3, 033111 (2021).

32. J. Berk, M. R. Foreman, Theory of Multiple Scattering Enhanced Single Particle Plasmonic Sensing. ACS Photonics 8, 2227–2233 (2021).

33. H. Lee, J. Berk, A. Webster, D. Kim, M. R. Foreman, Label-free detection of single nanoparticles with disordered nanoisland surface plasmon sensor. Nanotechnology 33, 165502 (2022).

34. L. Xiang, K. Chen, R. Yan, W. Li, K. Xu, Single-molecule displacement mapping unveils nanoscale heterogeneities in intracellular diffusivity. Nature Methods 17, 524–530 (2020).

35. J. A. Howarter, J. P. Youngblood, Optimization of Silica Silanization by 3-Aminopropyltriethoxysilane. Langmuir 22, 11142–11147 (2006).

