## Supplementary information for "Single-Objective Evanescent Scattering Microscopy for Imaging Single Proteins and Binding Kinetics"

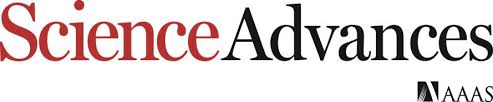

Supplementary Materials for

***Single-Objective Evanescent Scattering Microscopy for Imaging Single Proteins and Binding Kinetics***

Pengfei Zhang, Rui Wang, Zijian Wan, Xinyu Zhou, Guangzhong Ma,

Jayeeta Kolay, Jiapei Jiang, Shaopeng Wang*

**This PDF file includes:**

Fig. S1. Optical setup for single-objective evanescent scattering microscopy.

Fig. S2. Surface roughness profiles of ITO surface.

Fig. S3. Evaluation of heating effect.

Fig. S4. Estimation of shot noise limit.

Section S1. Theoretical prediction of signal-to-noise ratio.

Section S2. Estimation of super resolution localization.

**Other Supplementary Materials for this manuscript include the following:**

Movie S1.

Dynamic binding of BSA to anti-BSA and negative control experiment of flowing IgA onto anti-BSA.

Movie S2.

Tracking of one IgA protein binding on the surface.

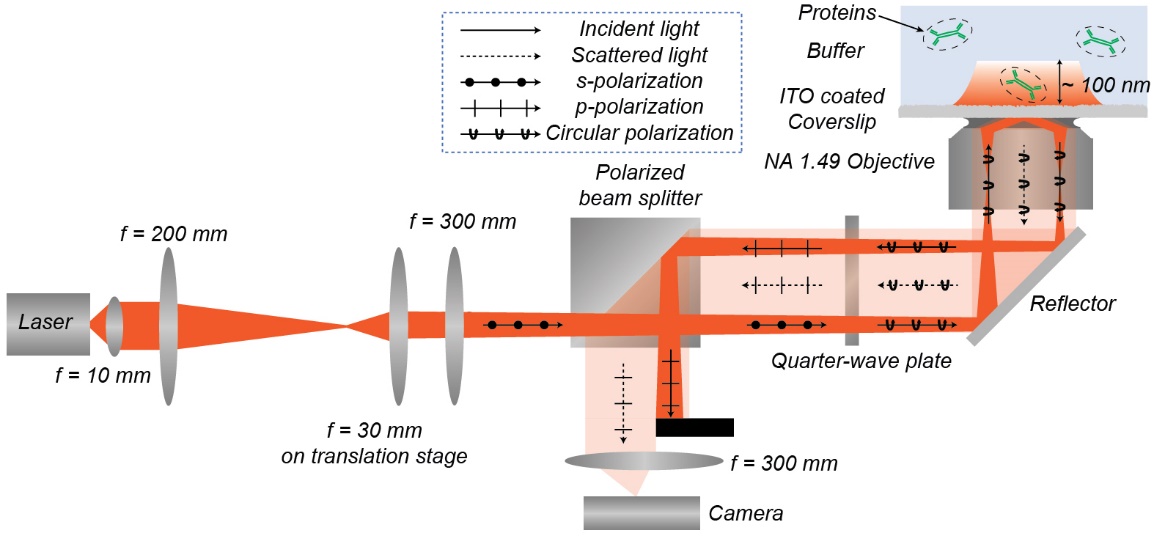

Fig. S1. Optical setup for single-objective evanescent scattering microscopy. Light from a laser diode with a center wavelength of 660 nm (L660P120, Thorlabs, Newton, NJ, US) is firstly collimated by a lens group with effective focal length of 10 mm. The lens group is constructed by two achromatic doublet lenses with focal length of 19 mm. The laser diode is fixed at a temperature-controlled mount (LDM56, Thorlabs), which is driven by a benchtop diode current controller (LDC205C, Thorlabs) and a temperature controller (TED200C, Thorlabs). Then, the light is conditioned again by a lens group constructed by two achromatic doublet lenses with focal length of 200 mm and 30 mm, respectively. The lens with focal length of 30 mm is placed in a manual three-dimensional translation stage (XR25P-K2, Thorlabs) for adjusting the incident angle beyond the incident angle. Then, the light is focused to the back focal plane of a ×60 objective (Olympus APO N 60x Oil TIRF, NA 1.49) by a tube lens with focal length of 300 mm. A combination of polarization beam splitter with quarter-wave plate is employed to separate the signal light from incident light. Refection beams is blocked by a M4 screw with a diameter of 4 mm. The evanescent waves scattered by the ITO surface and proteins are collected by a camera (MQ003MG-CM, XIMEA).

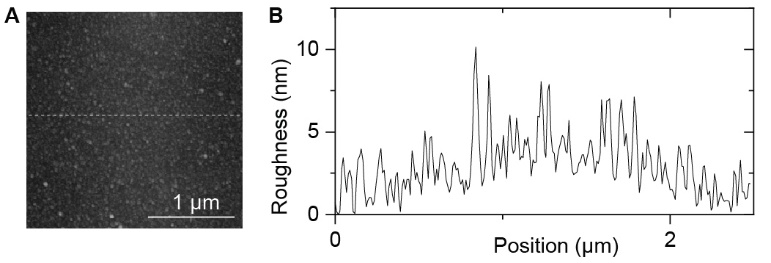

Fig. S2. Surface roughness profiles of ITO surface. (A) The surface of ITO coated coverslips was examined by Atomic Force Microscopy (Agilent 7500 AFM), showing islands of variable sizes. (B) Surface roughness profiles of marked line in (A).

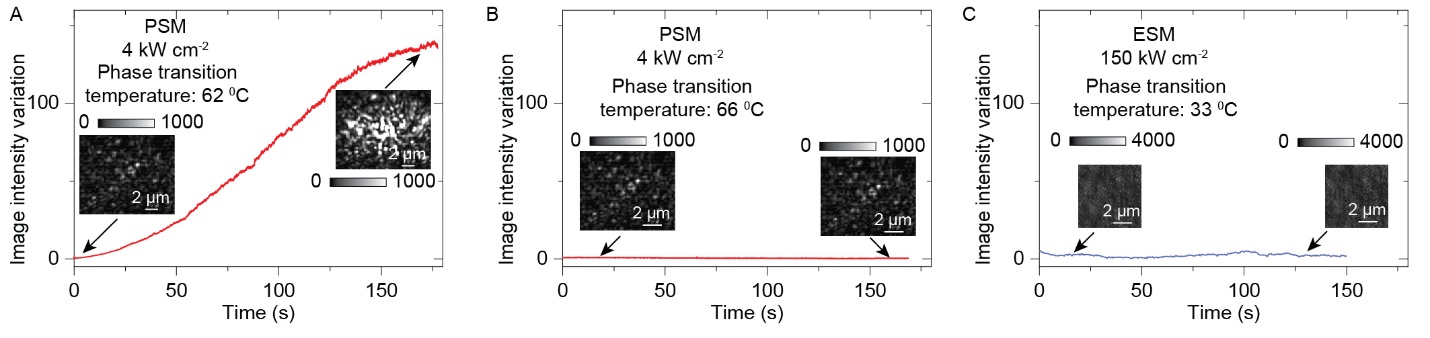

Fig. S3. Evaluation of heating effect. Some polymer solutions will produce large vesicles when solution temperature is higher than the phase transition temperature (J. Phys. Chem. B 2007, 111, 6, 1262–1270). The 0.236 g/L polyethylene oxide (PEO) dissolved in 300 mM NaH_2_PO_4_ and 240 mM NaH_2_PO_4_ aqueous solutions have the phase transition temperature at 62 °C and 66 °C, respectively. Phase transition was observed when flowing PEO in 300 mM NaH_2_PO_4_ solution onto the gold film surface of PSM (Fig. S3A), and not observed when flowing PEO in 240 mM NaH_2_PO_4_ solution (Fig. S3B). This indicates the PSM sensor surface temperature is between 62°C and 66 °C under the incident intensity of 4 kW cm^-2^. This is close to the upper-temperature limit for protein detection (Journal of Pharmaceutical and Biomedical Analysis 2020, 189, 113399), so the PSM usually uses 3 kW cm^-2^ as maximum incident intensity. In contrast, phase transition was not observed on the ITO surface of single-objective ESM under the incident intensity of 150 kW cm-2 when flowing 1 g/L cellulose in 300mM Na_2_HPO_4_ aqueous solution with a phase transition temperature of 33 °C. This indicates a much smaller heating effect on the ITO surface than the gold surface, thus allowing the single-objective ESM to employ the incident intensity of 150 kW cm^-2^, which is ~50 times higher incident intensity than reported value for PSM (Nat Methods 17, 1010–1017 (2020)). Camera exposure time is 0.1 ms for PSM, and 0.5 ms for ESM. A detailed study of the heating effect will be published in an upcoming article.

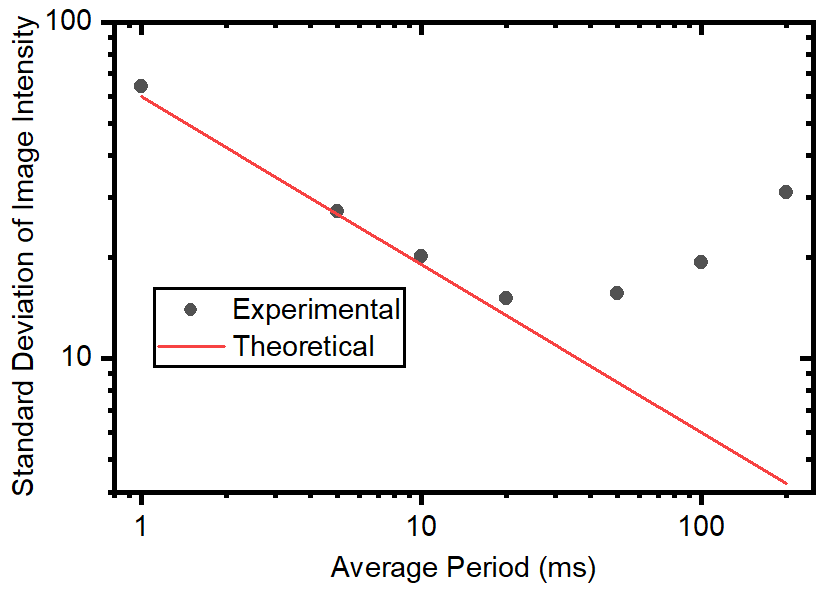

Fig. S4. Estimation of shot noise limit. Comparison of theoretical shot noise and experimental noise with different average periods, confirming that the shot noise is dominant for the average period below 20 ms. Incident light intensity, 150 kW cm^-2^. Exposure time, 0.5 ms. Detailed protocols have been described before (Nat Methods 17, 1010–1017 (2020)).

**Section S1. Theoretical prediction of signal-to-noise ratio**

The Rayleigh scattering model can also be used to estimate the system signal-to-noise ratio (SNR) theoretically. For the ESM measurement of single proteins, the image intensity *I* was determined by the interference between evanescent wave scattered by the surface roughness *E_b_* and protein *E_s_* as

$$I=\left| E_{b} \right|^{2}+2\left| E_{b} \right|\left| E_{s} \right|cos\beta+\left| E_{s} \right|^{2}, (S1.1)$$

where *β* is phase difference between *E_b_* and *E_s_*.

In the shot noise dominant system, the SNR is determined by the shot noise, which can be shown as

$$SNR=\frac{2\left| E_{b} \right|\left| E_{s} \right|cos\beta}{\sqrt{\left| E_{b} \right|^{2}+2\left| E_{b} \right|\left| E_{s} \right|cos\beta+\left| E_{s} \right|^{2}}}\approx\frac{2\left| E_{b} \right|\left| E_{s} \right|cos\beta}{\sqrt{\left| E_{b} \right|^{2}}}=2\left| E_{s} \right|cos\beta, (S1.2)$$

The phase difference *β* is about zero in ESM because of the short distance between scattering sites of surface roughness and analyte binding positions (ACS Photonics 8, 2227-2233 (2021)). The equation shows that the SNR is mainly determined by the scattering amplitude detected by the camera in the shot noise dominant system.

To estimate the theoretical SNR limit for the ESM system, the total Rayleigh scattering intensity *I_total_* of single protein is firstly estimated by

$$I_{total}=\frac{\frac{2\pi^{5}}{3}\times\frac{d^{6}}{\left( \frac{\lambda}{n_{m}} \right)^{4}}\times\left( \frac{\left( \frac{n_{s}}{n_{m}} \right)^{2}-1}{\left( \frac{n_{s}}{n_{m}} \right)^{2}+2} \right)^{2}}{A}\times P\times A\times t, (S1.3)$$

where *n_s_* and *n_m_* are the refractive indices of analyte and medium, *λ* is the incident wavelength, *d* is the analyte diameter, *P* is the incident light intensity, *t* is the average period. For the ESM used in this study, *n_s_* = 1.48, *n_m_* = 1.33, *λ* = 660 nm, and *t* = 0.02 s. Considering the single photon energy of ~1.2398/(0.66 µm) eV and the 5 x intensity enhancement of evanescent field, the total scattering intensity of one object in the ESM system can be estimated as

$$I_{total}={6\times10}^{-5}\times\left( d\left( nm \right) \right)^{6}\times P\left( kW{cm}^{-2} \right) photons. (S1.4)$$

The objective collects the scattering photons in perpendicular to the propagation direction of the evanescent wave, and the collection efficiency can be calculated with the equation in a spherical coordinate system of

$$\frac{I_{collection}}{I_{total}}=\frac{\int_{\theta_{1}}^{\theta_{2}} \int_{\varphi_{1}}^{\varphi_{2}} \frac{\left( 1+{cos}^{2}\theta\right)}{R^{2}}R^{2}sin\theta d\varphi d\theta}{\int_{0}^{\pi} \int_{0}^{2\pi} \frac{\left( 1+{cos}^{2}\theta\right)}{R^{2}}R^{2}sin\theta d\varphi d\theta}. (S1.5)$$

where $\theta$ and $\varphi$ are the polar angle and azimuthal angle, respectively. The objective collection angle for the ESM can be calculated by

$$\vartheta=arcsin\left( NA \right), (S1.6)$$

where *NA* is the objective numerical aperture. For the objective with NA of 1.49, the collection efficiency is calculated to be ~43 %. Considering the effect of bloker, the collection efficiency can be estimated to be ~36%.

For the detection of BSA with diameter of 7.4 nm under incident intensity of 150 kW cm^-2^ and average period of 20 ms, the total number of scattering photons is ~1480 based on equation (S1.4), and the number of photons collected by the objective is ~500 based on equations (S1.5) and (S1.6). Thus, the theoretical limit of SNR is ~ 45 under perfect conditions based on equation (S1.2). The transimission ratio of the imaging objective and tube lens in the ESM is measured to be ~70%, and the quantum efficiency of XIEMA MQ003 camera is ~48% for incident wavelength of 660 nm, so ~170 photoelectrons actually contribute to the sensor output. Considering of ~40% higher noise brought by the differential processing, the shot noise limited SNR of our setup for the BSA detection with ESM employing incident intensity of 120 kW cm^-2^ and average period of 20 ms is ~ 18, agreeing with the experimental results.

**Section S2. Estimation of super resolution localization**

For single particle tracking, the tracking precision in one dimension can be theoretically described by

$$\begin{aligned} \sigma_{\mu i}=\sqrt{\frac{s_{i}^{2}}{N}+\frac{a^{2}}{12N}+\frac{8\pi s_{i}^{4}b^{2}}{a^{2}N^{2}}},\#\left( S2.1 \right) \end{aligned}$$

where 𝜎_𝜇𝑖_ is the standard deviation of localization positions at *i*-th dimension, *N* is the photo number, *a* is the pixel size/magnification, *s* is the standard deviation of Gaussian distribution, and *b* is standard deviation of camera output in dark (Science, 300, 2061-2065 (2003)). The photon number *N* was used to estimate the shot noise limited signal to noise ratio (SNR), which is equal to the square root of photon number. For interference detection, the photon number *N* can be represented by the square of the shot noise limited SNR (Nat Commun 5, 4495 (2014)). The full width at half maximum (FWHM) can be estimated for optical imaging by dividing the incident wavelength by two times of imaging objective numerical aperture (NA), and the standard deviation can be estimated by dividing the FWHM by 2.35. Thus, the *s* can be estimated by

$$\begin{aligned} s_{i}=\frac{\lambda}{2\times2.35\times NA}.\#\left( S2.2 \right) \end{aligned}$$

The second and third terms in S2.1 are usually much smaller than the first term for interference measurement. Thus, the equation S2.1 can be simplified as

$$\begin{aligned} \sigma_{\mu i}\approx\frac{\lambda}{2\times2.35\times NA\times SNR}.\#\left( N3.4 \right) \end{aligned}$$

For the IgA measured under incident intensity of 150 kW cm^-2^ and average period of 10 ms, the SNR can be estimated to be ~70. The objective has NA of 1.49, and ~1.0 in horizontal, and vertical directions, respectively. Thus, the theoretical localization precision for IgA measured in Fig. 4C can be estimated to be 1.3 nm, and 2.0 nm in horizontal, and vertical directions, respectively.
